# Sodium channel inhibitors alter the progress of tangle development in a mouse model of dementia

**DOI:** 10.1101/2024.08.26.609302

**Authors:** Chloe M. Hall, Martha Roberts, Roshni A. Desai, Damian M. Cummings, Jamie Bilsland, Paul Whiting, Kenneth J. Smith, Frances A Edwards

## Abstract

Sodium channel inhibitors have been reported to protect against a range of neuroinflammatory and neurodegenerative diseases. Here the effect of chronic administration of two Na^+^ channel inhibitors with different mechanisms of action, phenytoin and GS967 are tested in mouse models of different stages of Alzheimer’s disease. Subtle changes in the distribution of plaque sizes were observed in *App*^*NLGF/NLGF*^ mouse at 3 months of age, after being fed control or drug-supplemented chow from weaning onwards, with phenytoin treatment resulting in a significant increase in the frequency of the smallest plaques and a decrease in large plaques. The later pathology of neurofibrillary tangles was studied, in old age, by supplementing the food of transgenic mice with a P301L mutation in Tau. Chronic administration of Na^+^ inhibitors from 15 months of age resulted in a decrease in the density of MC1-positive neurofibrillary tangles, possibly due to effects on microglial Na^+^ channels. The density of microglial cells was strongly correlated with the density of neurofibrillary tangles but only in mice treated with the Na^+^ inhibitors.

## INTRODUCTION

Sodium channel inhibitors have emerged as promising protective agents in various types of neuroinflammatory and neurodegenerative diseases. They have shown efficacy in diverse animal models including, but not limited to, experimental autoimmune encephalomyelitis (EAE; Lo *et al*., 2003; Bechtold *et al*., 2004; Bechtold *et al*., 2005; Bechtold *et al*., 2006; Morsali *et al*., 2013) and experimental Parkinson’s disease (Sadeghian *et al*., 2016). Moreover, clinical trials suggest promise for treating multiple sclerosis (Kapoor *et al*., 2010; Gnanapavan *et al*., 2013), optic neuritis (Raftopoulos *et al*., 2016) and Alzheimer’s disease (Golmohammadi *et al*., 2024). These studies have tested a range of drugs that have the common property of blocking voltage-gated sodium channels, including carbamazepine (Al-Izki *et al*., 2014), flecainide (Bechtold *et al*., 2004; Bechtold *et al*., 2005; Morsali *et al*., 2013), lamotrigine (Bechtold *et al*., 2006; Kapoor *et al*., 2010; Gnanapavan *et al*., 2013), oxcarbazepine (Al-Izki *et al*., 2014), phenytoin (Lo *et al*., 2003), riluzole (Simard *et al*., 2012; Wilson & Fehlings, 2014; Golmohammadi *et al*., 2024) and safinamide (Sadeghian *et al*., 2016).

Importantly, while it is accepted that all drugs have some off-target effects, the above compounds have the marked advantage of being safe for chronic administration in humans, with relatively benign side-effects.

Individuals with Alzheimer’s disease have a higher incidence of epilepsy compared to their age-matched counterparts without the disease (Bell *et al*., 2011). As a result, various drugs have been tested on transgenic mouse models of Alzheimer’s disease in relation to seizure activity, with a range of effects (Ziyatdinova *et al*., 2011; Sanchez *et al*., 2012; Verret *et al*., 2012). On the other hand, it should be noted, in the case of treating epilepsy with anticonvulsants, there is little evidence that this prevents development of Alzheimer’s disease (Carter *et al*., 2007). However, this may be because epilepsy itself increases the risk of Alzheimer’s disease. Whether decreasing Na^+^ channel activity would alter other aspects of the progression of the disease, in the absence of epilepsy, has not been tested.

Amyloidβ, which is released in an activity-dependent manner at synapses, increases glutamate release probability (Abramov *et al*., 2009). This observation implies a feed forward loop that might lead to ever increasing levels of Amyloidβ (Cirrito *et al*., 2008; Abramov *et al*., 2009). Therefore, disrupting this cycle may offer therapeutic advantages. Moreover, increased glutamate release has been described in a range of mouse models of Alzheimer’s disease, even before plaques become evident (Busche *et al*., 2012; Cummings *et al*., 2015; Benitez *et al*., 2021). Other evidence for hyperactivity has also been reported and this activity has been shown to be reversed by Na^+^ channel inhibitors (Ciccone *et al*., 2019). Notably, a recent study in humans has associated such hyperactivity with the development of neurofibrillary tangles (Giorgio *et al*., 2024). However, the effects of modulating such activity on the development of pathology has not been investigated.

We have therefore used two approaches to inhibit Na^+^ channel activity through the chronic administration of either: 1. phenytoin, a long established anticonvulsant drug that blocks high-frequency action potentials that underlie the abnormal activity associated with seizures, while sparing normal brain activity (Yaari *et al*., 1986); or 2. GS967, a non-inactivating sodium channel blocker; that preferentially blocks the Na^+^ channel late current rather than the peak current (Anderson *et al*., 2014). To investigate potential benefits of these Na^+^ channel inhibitors, we have used two mouse models to examine effects on different disease pathologies. Firstly, to address the earliest stages of Alzheimer’s disease, we tested the effects of administration of Na^+^ channel inhibitors on plaques in the APPKI mouse *App*^*NLGF/NLGF*^ (Saito *et al*., 2014). We hypothesised that disrupting the feed forward loop of activity-dependent release of Amyloidβ might be beneficial. Secondly, as a proxy for the later clinical stages of tauopathies such as Alzheimer’s disease, we observed the development of tau tangles in TauD35 P301L mice (Joel *et al*., 2018), a model of frontotemporal dementia with parkinsonism (FTDP). We thus investigated whether interfering with the observed association of hyperactivity with Tau pathology (Giorgio *et al*., 2024) could alter the development of tangles. Neither of these mouse models has been reported to feature detectable epileptic episodes. Hence we are not investigating the role of epilepsy in Alzheimer’s disease but rather the role of hyperactivity and whether decreasing this activity could be a potential therapeutic strategy.

Na^+^ channel antagonists have also been reported to have potent effects in suppressing the pro-inflammatory activation of microglia within the brain (Craner *et al*., 2005; Sadeghian *et al*., 2016), which is especially important given the growing evidence regarding microglial activation and proliferation in Alzheimer’s disease. In this context, we have also investigated whether the drugs influence the microglial proliferation that has previously shown to be in response to proximity to plaques and, to a lesser degree, tangles (Matarin *et al*., 2015; Chen *et al*., 2020; Wood *et al*., 2022).

## MATERIALS AND METHODS

### Animal models

All mice were housed in groups of 2-5 animals in enriched environments, or occasionally single housed for a maximum of 24 hours at the Biological Services Unit of University College London. Access to food and water was *ad libitum* and mice were kept on a 12-hour light/dark cycle. All procedures were performed in accordance with the United Kingdom Animals (Scientific Procedures) Act 1986.

### APP knock in mice

Male homozygous *App*^*NL-G-F/NL-G-F*^ knock-in mice (NLGF) aged 3-3.5 months were used. These mice have a humanised Amyloidβ sequence within *App* and harbour Swedish (KM670/671NL), Beyreuther/Iberian (I716F) and Arctic (E694G) mutations (Saito *et al*., 2014). In the hippocampi of NLGF mice Amyloidβ plaques are first detectable at around 2 months (Saito *et al*., 2014). For further characterisation of these mice see (Benitez *et al*., 2021).

### TauP301L mice

In a separate series of experiments to study the development of Tau pathology the mice used were transgenic for human Tau with a P301L mutation driven by the CaMKII promoter. This mutation causes frontotemporal dementia with parkinsonism linked to chromosome 17. These mice were developed by GlaxoSmithKline and it is now known that there are mice with two different transgene copy numbers (Joel *et al*., 2018). The line with high copy number and higher Tau levels were used for this study.

Tau tangle-like pathology first appears at 8 months of age in these mice, and they reach severe neurodegeneration with a hunched, piloerect, and akinesic phenotype at about 17 months of age. According to Home Office requirements, mice were closely monitored and killed when the neurodegenerative phenotype became evident. The brain was immediately dissected out, and one hemisphere of the brain was drop fixed in 4% paraformaldehyde in phosphate-buffered saline (PBS) for 24 hours and then stored in 30% sucrose, 0.03% NaN_3_ in PBS at 4°C.

### Chronic Administration of Drugs

Two Na^+^ channel inhibitors were investigated in this experiment: phenytoin and GS967. The drugs were administered orally through supplementation in chow. For investigation of early plaque development in the *App*^*NL-G-F/NL-G-F*^ mice supplemented chow was available *ad libitum* from weaning at 3 weeks of age. For investigation of tau tangles in TauP301L mice supplemented chow was available *ad libitum* from 15 months of age. Age matched mice of both genotypes were administered the same chow, without drugs as controls. All other conditions remained the same. Drug doses used were 336 mg/kg of diet for phenytoin and 8 mg/kg of diet for GS967. Drug dosage was determined with reference to previous studies (Craner *et al*., 2005; Anderson *et al*., 2014).

### Immunohistochemistry

One brain hemisphere from each animal was drop fixed in 4% paraformaldehyde in PBS for 24 hours and then stored in 30% sucrose, 0.03% NaN_3_ in PBS at 4°C. Hemispheres were cut to optimise sections transverse to the hippocampus and embedded in OCT compound (BDH, Poole, UK), frozen, and stored until use at -80°C. Frozen cryostat sections (15 µm) were mounted onto Superfrost Plus slides (VWR International, UK) and kept at -80°C until staining. Brain sections were labelled for MC1 using immunohistochemistry. The primary antibody and the corresponding secondary antibody are given in Table 1. Slides were air dried overnight before any histology. Sections were rehydrated by washing in PBS buffer. Non-specific binding was blocked using 5% goat serum in 0.1 M PBS for at least 30 mins. Sections were incubated with primary antibody overnight at 4°C. Primary antibodies were detected using a secondary antibody fused to AlexaFluor488 or AlexaFluor594 at a 1:500 dilution or by diaminobenzidine staining using standard methods (Table 1). Incubation of secondary antibodies was for two hours at room temperature. Three PBS washes were performed between each step. Sections were coverslipped with Vectashield (Vector Laboratories) containing DAPI.

**Table 1.** Details of antibodies used.

| Primary Antibody | Antigen | Dilution | Secondary Antibody | Chromogen/Fluorophore |
| --- | --- | --- | --- | --- |
| Mouse anti-MC1 (Gift from Peter Davies) | Misfolded human Tau | 1:200 | Goat-anti mouse IgG | AlexaFluor 594 |
| Rabbit anti-IBA1 (Wako) | General microglial marker | 1:500 | Goat-anti rabbit IgG | Diaminobenzidine |
| Rabbit anti-Aβ42 (ThermoFisher) | Aβ1-42 | 1:300 | Donkey anti-rabbit IgG | AlexaFluor 488 |

### Imaging

Slides were imaged with an EVOS Auto FL Microscope (Life Technologies) using a x20 objective including the whole hippocampus and some surrounding cortex.

### Statistical analysis

Quantitative data are presented as mean ± standard error of the mean (SEM) throughout. Sample sizes are referred to as number of animals for each experimental group. Outliers were classified as data more than two standard deviations away from the mean and were excluded from further analysis. Two-way analysis of variance (ANOVA) and unpaired t-tests were used to assess significance between groups. Tukey post-hoc tests were used throughout. Differences were considered statistically significant when p<0.05. All statistical tests and plots were performed using Prism v10 (Graphpad).

## RESULTS

### Effects of chronic treatment with Na^+^ channel inhibitors on mice with plaques

#### Na^+^ channel inhibitors affect the early development of plaques in App^NL-G-F/NL-G-F^ mice

To assess the effects of inhibiting Na^+^ channels on the early deposition of plaques, *App*^*NL-G-F/NL-G-F*^ *mice* were fed either normal mouse chow or chow containing 340 mg/kg of diet phenytoin or 8 mg/kg of diet GS967 from weaning (from 3 weeks of age, before plaques are evident) until use in experiments at 3 – 3.5 months of age. The weights of 3 mice from each treatment group were measured regularly. Growth rates were not significantly different between groups. Weight gain over a 6-week period was: control diet, 10.2 ± 0.9 g; Phenytoin, 11.6 ± 1.2 g; GS967, 10.6 ± 0.9 g.

Plaques were still fairly sparse at the time of the analysis. Although there was no significant main effect of treatment on plaque density (data not shown), there was a strong trend for treatment to affect median plaque area (one-way ANOVA, treatment p = 0.055 with a significant effect of phenytoin, p = 0.035; Fig. 1A-C). The pooled distribution of plaque sizes was significantly affected by the chronic ingestion of phenytoin with the distribution being skewed towards a higher proportion of very small plaques (Fig 1b) compared to control mice, with control mice showing the largest plaques. In control mice, a small proportion of large plaques (180 – 1306 µm^2^) were seen in 4 of the 5 control mice, whereas only 1 mouse in each of the treatment groups showed one or two plaques in this range (192 – 216 µm^2^). (Control n = 5; treatments n = 4 animals).

**Figure 1.**
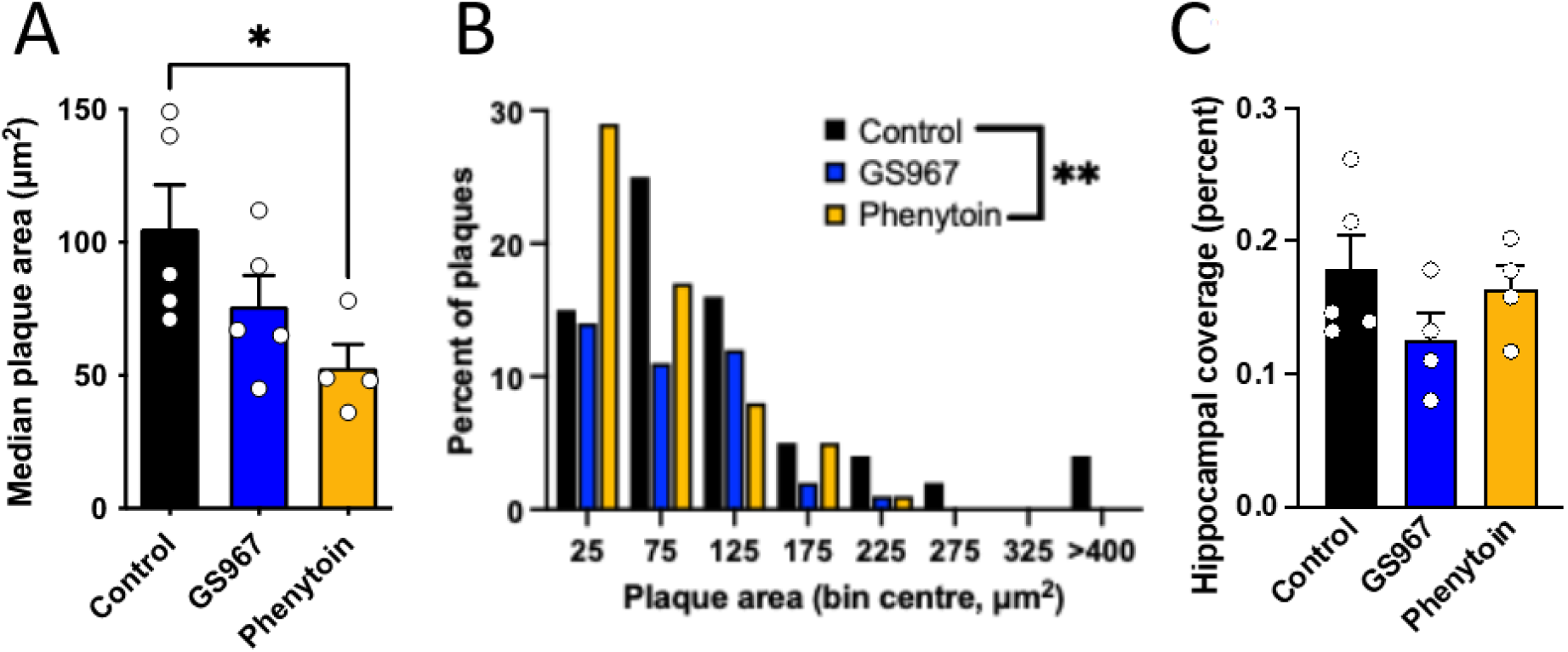
Treatment with phenytoin results in a higher proportion of small plaques and prevent the occurrence of the largest plaques. A. Mean area of individual plaques; B. Frequency distribution of plaque sizes; C. Percentage of hippocampus covered by plaque. Individual points represent individual mice; columns represent mean ± S.E.M. Asterisks represent posthoc analysis *p < 0.05, **p < 0.01

### Effects of chronic treatment with Na^+^ channel inhibitors on mice with neurofibrillary tangles

#### Chronic treatment with Na^+^ channel inhibitors prevented loss of brain weight but did not improve life span in TauP301L mice

The cognitive deficits of Alzheimer’s disease only become apparent once neurofibrillary tangles begin to build up in the hippocampus (Nelson *et al*., 2012; Edwards, 2019). As treatment to prevent the progression of disease would be more likely to be applicable to these later stages of disease, once symptoms became apparent, we concentrated our study on the development of neurofibrillary tangles. TauP301L mice were fed GS967 or phenytoin or control diet from 15 months old until the neurodegenerative phenotype was observed, at which point animals were culled. At this age mouse growth has plateaued and there was no difference over a 6-week period in the weight of mice on different diets. The neurodegenerative phenotype was visually evident as hunched posture, piloerection and complete immobility. As a rough assessment of the degree of neurodegeneration the fixed brains of the mice were blotted dry and weighed before sectioning.

Mice reached the neurodegenerative phenotype between 16.4 and 19.6 months. There was no significant difference between the treatment groups (age in months): controls, 18.21 ± 0.44, n = 7; GS967, 16.93 ± 0.27, n = 4; phenytoin, 17.93 ± 0.17, n = 3; Fig. 2A)). The hemisphere of the brain to be used for immunohistochemistry was weighed at the endpoint. On the control diet, TauP301L mice were found to have a significantly lower brain weight than WT mice but this decrease was prevented by both drugs (Fig. 2B,C).

**Figure 2.**
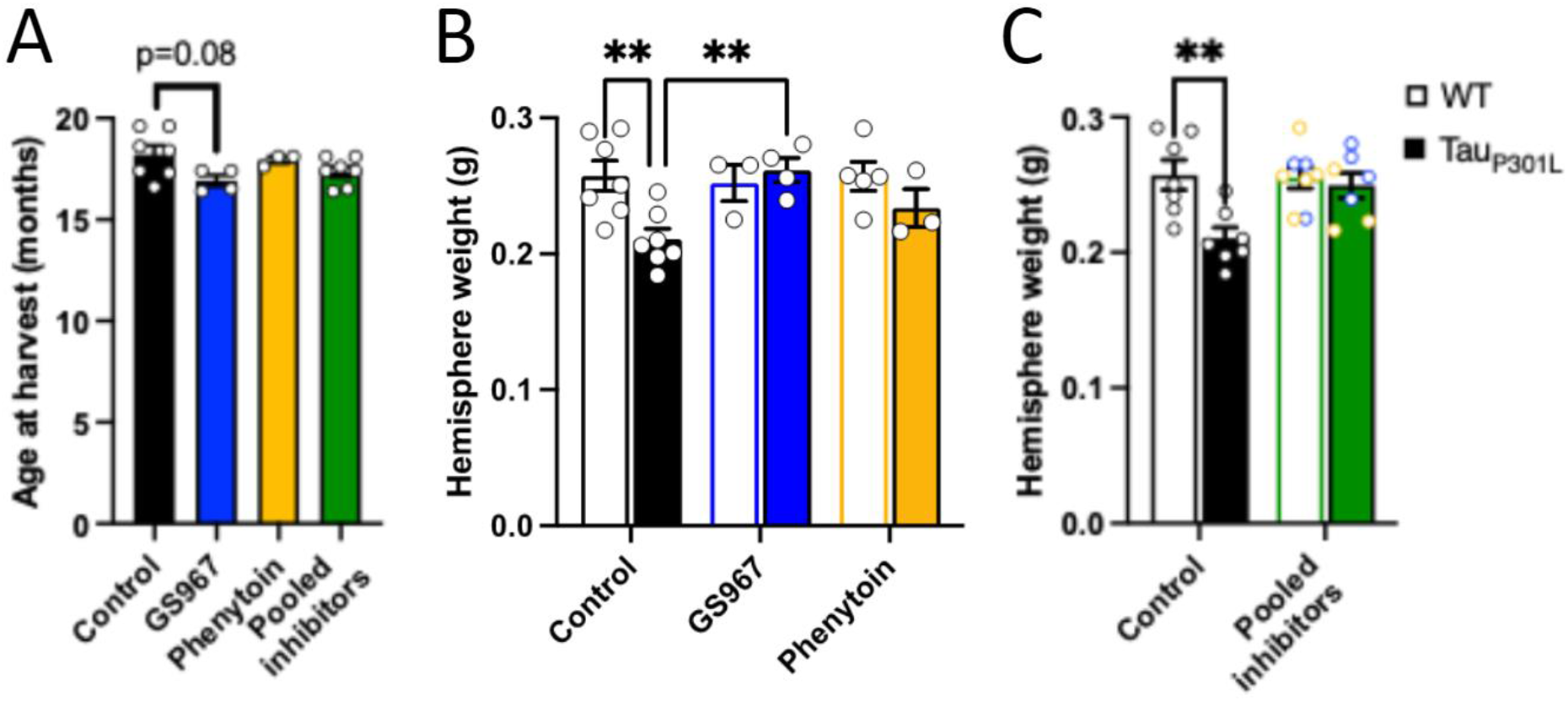
A. Treatment with Na^+^ channel inhibitors did not significantly change the age of reaching the neurodegenerative phenotype. B TauP301L mice had significantly lower brain weights than WT mice but this difference was lost after treatment with either drug. C. Pooling the brain weights of mice treated with inhibitors shows no significant difference between brain weights of WT and TauP301L mice. Asterisks represent posthoc analysis ** p < 0.01

#### Chronic treatment with Na^+^ channel inhibitors decreased tangle load in TauP301L mice

To assess the effect of the two Na^+^ channel inhibitors on the development of neurofibrillary tangles, sections were labelled with an MC1 antibody which labels paired helical filaments (Jicha *et al*., 1997). Positive labelling that clearly surrounded a DAPI nucleus was considered indicative of the presence of a neurofibrillary tangle. Tangles were manually counted per Area of Interest (AOI, 240 µm x 400 µm) in the cell body layer of each of the primary hippocampal regions (dentate gyrus, CA3 and CA1) with AOIs from 2 hippocampal sections per animal averaged to give n = 1. A trend was seen across all regions to show a decrease in MC1 positive cell bodies when the two drug treatments were analysed separately (2-way ANOVA main effect treatment, p = 0.07). Interestingly there was a significant difference between hippocampal areas, with the dentate gyrus showing significantly more tangles than the other regions (main effect of region, p = 0.02). As sample sizes for the treatment groups were small, the two treatment groups were pooled. Pooling the treatment groups revealed a significant decrease in the density of MC1 positive cell bodies (p < 0.02; Fig. 3).

**Figure 3.**
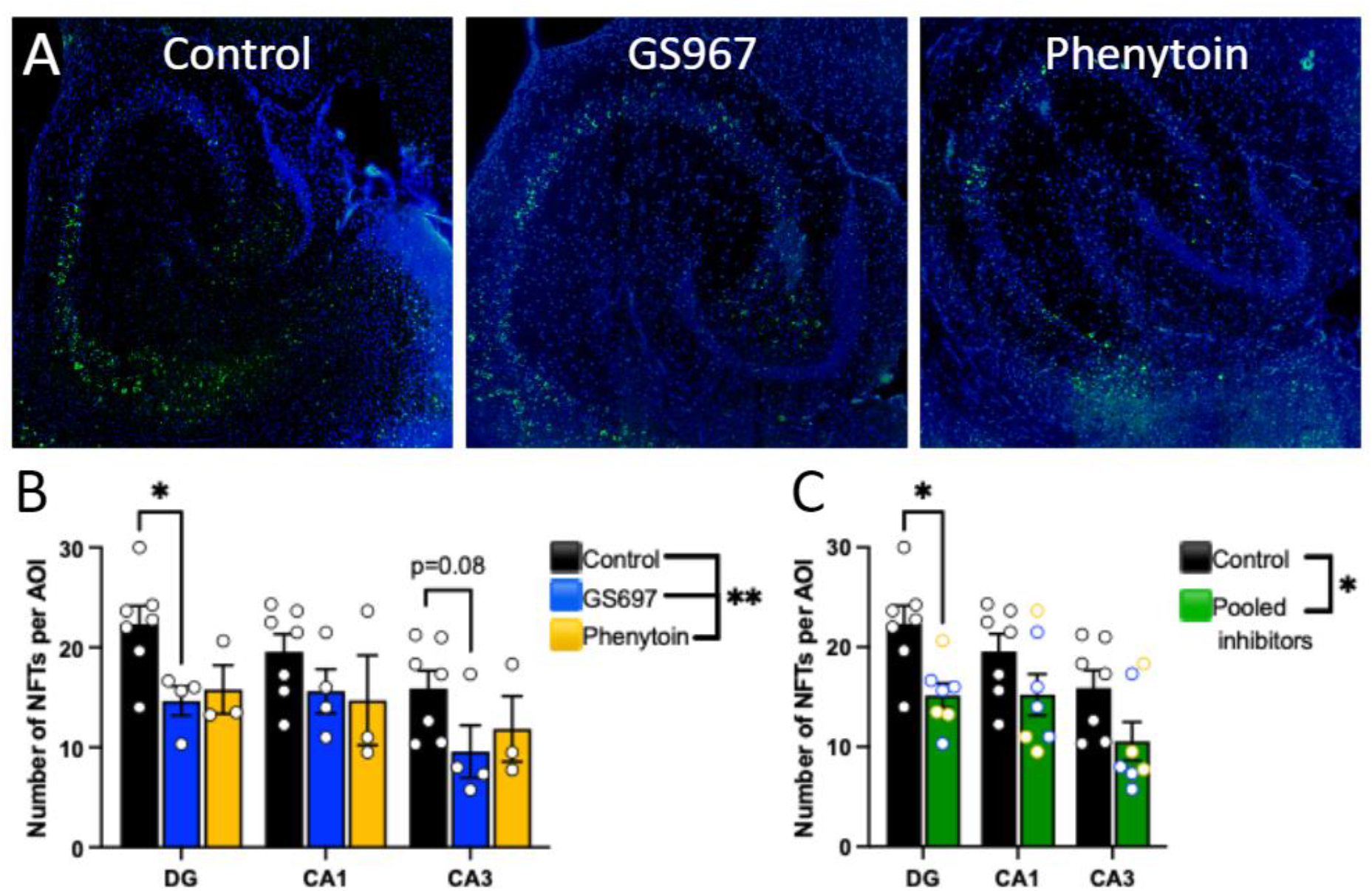
Na^+^ channel inhibitors decrease the density of neurofibrillary tangles. A. Images of DAPI-stained (blue) and MC1 (green) positive neurofibrillary tangles in aged hTauP301L mice, fed with the treatment indicated. B,C. Counts of neurofibrillary tangles in AOIs (400µm x 240µm) in the cell body layer of different regions of the hippocampus. B. Effects of individual treatments do not reach statistical significance. C. Pooling the results from the two Na^+^ channel inhibitors shows a significant decrease in tangle load. 2-way analysis of variance: main effects of region (p = .0004) and treatment (p = 0.02). Asterisks represent posthoc analysis * p < 0.05.

### Treatment with Na^+^ channel inhibitors did not affect microgliosis in TauP301L mice

#### Microglia density increased in TauP301L mice compared to WT mice but this difference was not affected by treatment with individual Na^+^ channel inhibitors

As microglia are clearly important in the progression of various forms of dementia we assessed the density of IBA1-positive microglia in WT versus TauP301L mice in the different layers of the CA1 region and in the dentate gyrus. In the TauP301L mice there is an increase in microglial number compared to WT in both CA1 region and in the dentate gyrus but no significant difference is seen with either drug treatment. Counts were made in different CA1 layers and in both CA1 and dentate gyrus with similar results in all regions (Fig. 4). As the trend for dentate gyrus was in the same direction for these two related treatments, we again pooled the data to assess whether there was an overall effect of Na^+^ channel inhibitors again revealing a significant effect of Na^+^ channel inhibitors, (p = 0.03; Fig. 4D). This suggests that the inhibition of Na^+^ channels decreases the response of the microglia in the TauP301L mice bringing it almost back to WT levels which could have important effects on disease progression.

**Figure 4.**
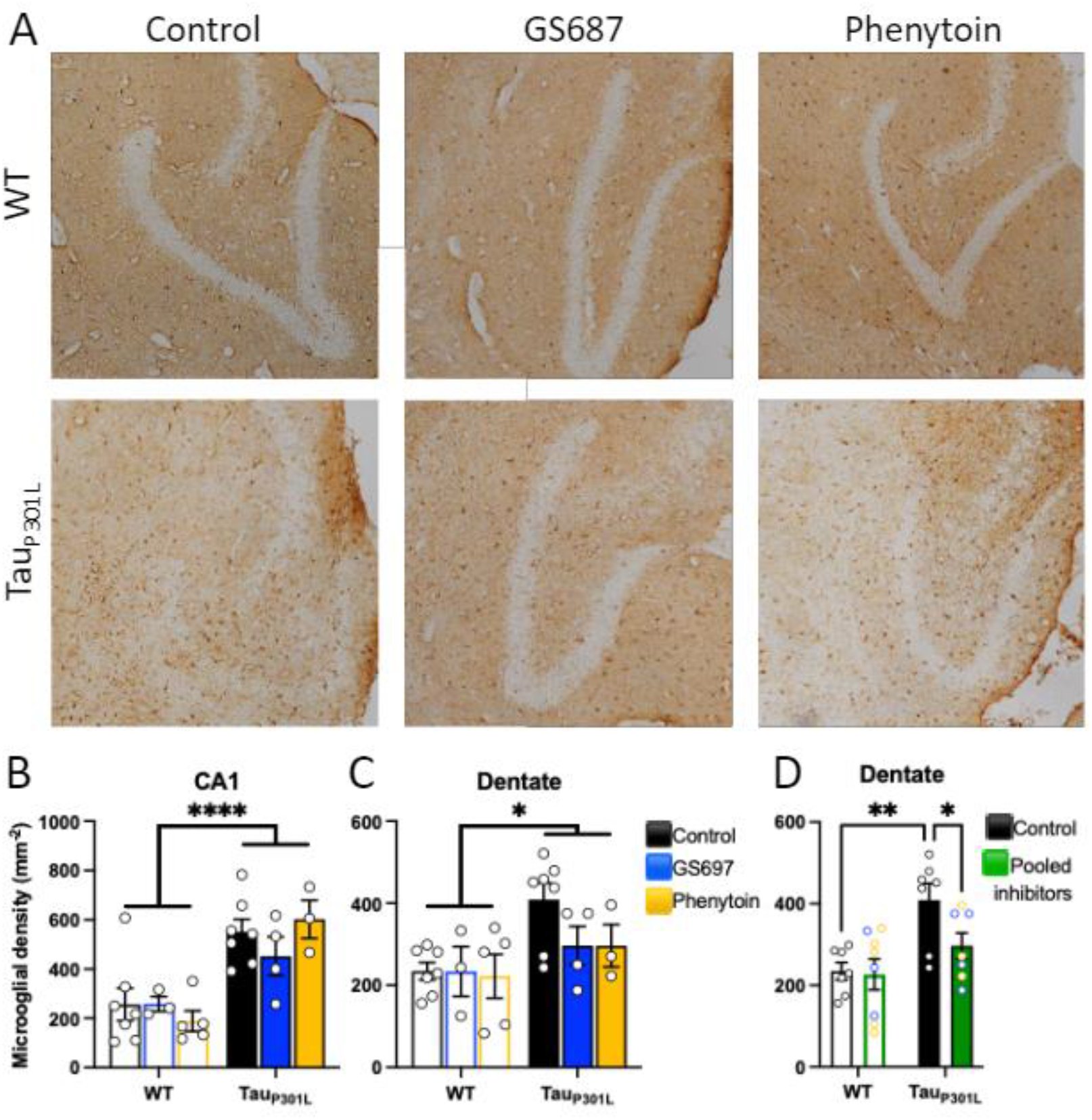
Treatment with Na^+^ channel inhibitors decreased microglial proliferation in the dentate gyrus of TauP301L mice A. Iba1 stained cells were counted in fixed tissue in different regions of the hippocampus. B-C. Results from CA1 and dentate regions respectively. D. Pooled data from the two treatment groups showed a significant effect of inhibiting Na^+^ channels. 3 sections per mouse were analysed and the results averaged. Individual data points refer to means for individual animals. Asterisks represent posthoc analysis* p < 0.05, ** p < 0.01, **** p < 0.0001.

Considering that the differences in both neurofibrillary tangle density and microglial density were in the same direction with the two treatments but showed considerable variation, we investigated whether the density of tau tangles correlated with the density of microglia in the same region. While a significant correlation was observed if all groups were pooled, the correlation was clearly driven by the treatment groups, with no correlation being observed in the control group alone (Fig. 5). Not surprisingly such correlations were not observed in the CA1 region where changes in these variables were less consistent.

**Figure 5.**
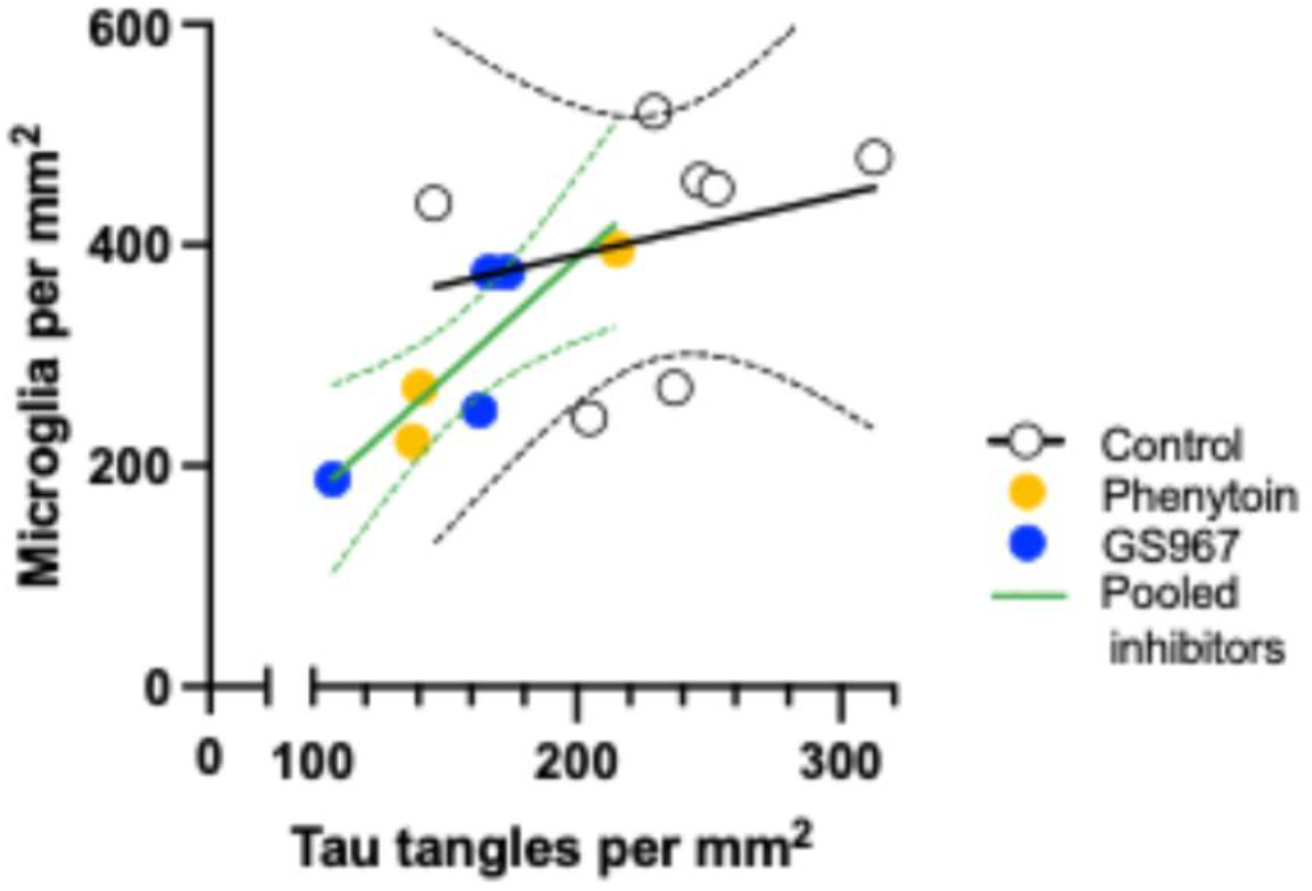
In dentate gyrus, the density of neurofibrillary tangles correlated with the density of microglia. Pooling data from all groups resulted in a significant correlation (Pearson correlation 0.64; p < 0.02). Correlations did not reach significant in individual treatment groups (n = 3-4) but if Na^+^ channel inhibitors were pooled, to compare equal sample sizes (n = 7) between pooled treatment and controls, a strong correlation was observed for pooled treatment groups (Pearson correlation r = 0.87; p = 0.01). The control group alone showed no significant correlation (Pearson correlation 0.25; p = 0.58). Control diet open black circles; GS967 filled blue circles; phenytoin filled yellow circles. Linear regression lines and 95% confidence intervals shown for control diet (black) and pooled treatments (green).

## DISCUSSION

Modulating Na^+^ currents with phenytoin or GS967 had subtle effects on both plaque deposition and the density of neurofibrillary tangles.

This study investigated the initial deposition of plaques in APPKI mice (*App*^*NLGF/NLGF*^) on the basis that the release of Amyloidβ is activity-dependent (Abramov *et al*., 2009) and thus administration of inhibitors of Na^+^ channels could influence the initial seeding of plaques. Small plaques are first detectable in the *App*^*NLGF/NLGF*^ from 2 months of age and so have only been developing for a short period by the age tested (3.5 months). However, a surprising spread of plaque sizes was evident in the mice on control diet. This heavily skewed distribution was truncated in the mice treated with Na^+^ channel inhibitors, suggesting that the ongoing growth of the plaque, after initial seeding, continues to be activity-dependent. Other parameters including the density and overall plaque coverage were not changed. However, it is important to note that plaque number and coverage at this age are relatively low and rapidly increasing which likely contributed to the wide variability of the measurement.

In contrast to the amyloid mice the transgenic TauP301L mice were studied at the end stage of neurodegeneration. A cautionary note should be added as to the validity of the model. This, like all the previous tauopathy models, is an overexpression model of mutated Tau. We have published evidence that the overexpression itself may be an important factor in the build-up of Tau tangles and in the age of neurodegeneration in these models (Joel *et al*., 2018). The amount of Tau that can bind to the microtubules may be related to activity and so this could interact with these treatments. Furthermore, this is a model for frontotemporal dementia with parkinsonism linked to chromosome-17, rather than Alzheimer’s disease; however, the effects of tau tangles are most likely common to many related neurodegenerative diseases.

There was a strong trend for a decrease in tangle load after treatment with phenytoin or GS967 and this reached significance when the treatment groups were pooled for analysis. Moreover, the decreased brain weight measured in the mice on control diet was prevented by the Na^+^ channel inhibitors although this did not translate to an increase in healthy life span. Indeed, all groups of Tau mice displayed the neurodegenerative phenotype at the same mean age independent of treatment. This would suggest that the tangle load or neurodegeneration as measured by brain weight were not the only driving forces for the neurodegenerative phenotype. Alternatively, these putatively protective changes were not sufficient to impact the phenotype.

One explanation for a decrease in neurofibrillary tangles in the treated mice could be that the dystrophic neurones containing hyperphosphorylated Tau were more efficiently removed by microglia under these conditions. However, this seems unlikely as, although there was a proliferation of microglia in the Tau mice compared to controls, the trend was for this proliferation to be decreased in treated mice. As loss-of-function mutations in some anti-inflammatory microglial genes, such as *TREM2*, increase the risk of Alzheimer’s disease (Guerreiro *et al*., 2013; Jonsson *et al*., 2013), it is likely that microglial activity slows the development of the disease at early stages, while at later stages they may cause increased neurodegeneration through release of proinflammatory cytokines. It is interesting to note that, when individual mice are compared, microglial density correlates strongly with neurofibrillary tangle load, but only in treated animals. The lack of correlation in the control group is consistent with previous observations in untreated mice that when mice with different tangle loads are compared, there is only a weak correlation with increased expression of microglial genes (Matarin *et al*., 2015). The strong correlation in the presence of Na^+^ channel inhibitors may suggest a role of microglial Na^+^ channels in this context. If microglial density were simply a response to tangle load then, although the tangle load might be influenced by decreased neuronal activity in treated mice, the correlation between tangle load and microglial density should not be changed by treatments. However, as the correlation is only seen in the treated mice, this suggests that a change in microglial activity may be influencing the tangle load. Indeed it has been reported that microglia express Nav1.6 Na^+^ channels and that Na^+^ channel inhibitors can reduce microglial activation (Black *et al*., 2009; Black & Waxman, 2012; Morsali *et al*., 2013; Sadeghian *et al*., 2016). This is a particularly interesting finding in the light of the likely protective effects of microglial activation around plaques in earlier stages of disease progression as evidenced by the increase in risk of developing Alzheimer’s disease when microglial response to plaques is defective such as in individuals with certain variants of the microglial *TREM2* gene (Guerreiro *et al*., 2013; Jonsson *et al*., 2013; Liu *et al*., 2020). It is thus surprising that interfering with microglial function would in this case be protective, with this difference, presumably relating to age and possibly specifically in relation to neurofibrillary tangles. Further functional studies would be needed to confirm the role of microglial Na^+^ channels in this context but it may confirm that treatments related to microglial activation could be valuable in various tauopathies. However, in Alzheimer’s disease this would relate specifically only to late stage disease with microglial activation having an important protective function in early disease (Edwards, 2019).

## REFERENCES

Abramov, E., Dolev, I., Fogel, H., Ciccotosto, G.D., Ruff, E. & Slutsky, I. (2009) Amyloid-beta as a positive endogenous regulator of release probability at hippocampal synapses. Nat Neurosci, 12, 1567–1576.

Al-Izki, S., Pryce, G., Hankey, D.J., Lidster, K., von Kutzleben, S.M., Browne, L., Clutterbuck, L., Posada, C., Edith Chan, A.W., Amor, S., Perkins, V., Gerritsen, W.H., Ummenthum, K., Peferoen-Baert, R., van der Valk, P., Montoya, A., Joel, S.P., Garthwaite, J., Giovannoni, G., Selwood, D.L. & Baker, D. (2014) Lesional-targeting of neuroprotection to the inflammatory penumbra in experimental multiple sclerosis. Brain, 137, 92–108.

Anderson, L.L., Thompson, C.H., Hawkins, N.A., Nath, R.D., Petersohn, A.A., Rajamani, S., Bush, W.S., Frankel, W.N., Vanoye, C.G., Kearney, J.A. & George, A.L., Jr. (2014) Antiepileptic activity of preferential inhibitors of persistent sodium current. Epilepsia, 55, 1274–1283.

Bechtold, D.A., Kapoor, R. & Smith, K.J. (2004) Axonal protection using flecainide in experimental autoimmune encephalomyelitis. Ann Neurol, 55, 607–616.

Bechtold, D.A., Miller, S.J., Dawson, A.C., Sun, Y., Kapoor, R., Berry, D. & Smith, K.J. (2006) Axonal protection achieved in a model of multiple sclerosis using lamotrigine. J Neurol, 253, 1542–1551.

Bechtold, D.A., Yue, X., Evans, R.M., Davies, M., Gregson, N.A. & Smith, K.J. (2005) Axonal protection in experimental autoimmune neuritis by the sodium channel blocking agent flecainide. Brain, 128, 18–28.

Bell, J.S., Lonnroos, E., Koivisto, A.M., Lavikainen, P., Laitinen, M.L., Soininen, H. & Hartikainen, S. (2011) Use of antiepileptic drugs among community-dwelling persons with Alzheimer’s disease in Finland. J Alzheimers Dis, 26, 231–237.

Benitez, D.P., Jiang, S., Wood, J., Wang, R., Hall, C.M., Peerboom, C., Wong, N., Stringer, K.M., Vitanova, K.S., Smith, V.C., Joshi, D., Saito, T., Saido, T.C., Hardy, J., Hanrieder, J., De Strooper, B., Salih, D.A., Tripathi, T., Edwards, F.A. & Cummings, D.M. (2021) Knock-in models related to Alzheimer’s disease: synaptic transmission, plaques and the role of microglia. Mol Neurodegener, 16, 47.

Black, J.A., Liu, S. & Waxman, S.G. (2009) Sodium channel activity modulates multiple functions in microglia. Glia, 57, 1072–1081.

Black, J.A. & Waxman, S.G. (2012) Sodium channels and microglial function. Exp Neurol, 234, 302–315.

Busche, M.A., Chen, X., Henning, H.A., Reichwald, J., Staufenbiel, M., Sakmann, B. & Konnerth, A. (2012) Critical role of soluble amyloid-β for early hippocampal hyperactivity in a mouse model of Alzheimer’s disease. Proc Natl Acad Sci U S A, 109, 8740–8745.

Calabresi, P., Centonze, D., Marfia, G.A., Pisani, A. & Bernardi, G. (1999) An in vitro electrophysiological study on the effects of phenytoin, lamotrigine and gabapentin on striatal neurons. Br J Pharmacol, 126, 689–696.

Carter, M.D., Weaver, D.F., Joudrey, H.R., Carter, A.O. & Rockwood, K. (2007) Epilepsy and antiepileptic drug use in elderly people as risk factors for dementia. J Neurol Sci, 252, 169–172.

Chen, W.T., Lu, A., Craessaerts, K., Pavie, B., Sala Frigerio, C., Corthout, N., Qian, X., Lalakova, J., Kuhnemund, M., Voytyuk, I., Wolfs, L., Mancuso, R., Salta, E., Balusu, S., Snellinx, A., Munck, S., Jurek, A., Fernandez Navarro, J., Saido, T.C., Huitinga, I., Lundeberg, J., Fiers, M. & De Strooper, B. (2020) Spatial Transcriptomics and In Situ Sequencing to Study Alzheimer’s Disease. Cell, 182, 976–991 e919.

Ciccone, R., Franco, C., Piccialli, I., Boscia, F., Casamassa, A., de Rosa, V., Cepparulo, P., Cataldi, M., Annunziato, L. & Pannaccione, A. (2019) Amyloid beta-Induced Upregulation of Na(v)1.6 Underlies Neuronal Hyperactivity in Tg2576 Alzheimer’s Disease Mouse Model. Sci Rep, 9, 13592.

Cirrito, J.R., Kang, J.E., Lee, J., Stewart, F.R., Verges, D.K., Silverio, L.M., Bu, G., Mennerick, S. & Holtzman, D.M. (2008) Endocytosis is required for synaptic activity-dependent release of amyloid-beta in vivo. Neuron, 58, 42–51.

Craner, M.J., Damarjian, T.G., Liu, S., Hains, B.C., Lo, A.C., Black, J.A., Newcombe, J., Cuzner, M.L. & Waxman, S.G. (2005) Sodium channels contribute to microglia/macrophage activation and function in EAE and MS. Glia, 49, 220–229.

Cummings, D.M., Liu, W., Portelius, E., Bayram, S., Yasvoina, M., Ho, S.H., Smits, H., Ali, S.S., Steinberg, R., Pegasiou, C.M., James, O.T., Matarin, M., Richardson, J.C., Zetterberg, H., Blennow, K., Hardy, J.A., Salih, D.A. & Edwards, F.A. (2015) First effects of rising amyloid-beta in transgenic mouse brain: synaptic transmission and gene expression. Brain, 138, 1992–2004.

Edwards, F.A. (2019) A Unifying Hypothesis for Alzheimer’s Disease: From Plaques to Neurodegeneration. Trends Neurosci, 42, 310–322.

Giorgio, J., Adams, J.N., Maass, A., Jagust, W.J. & Breakspear, M. (2024) Amyloid induced hyperexcitability in default mode network drives medial temporal hyperactivity and early tau accumulation. Neuron, 112, 676–686 e674.

Gnanapavan, S., Grant, D., Morant, S., Furby, J., Hayton, T., Teunissen, C.E., Leoni, V., Marta, M., Brenner, R., Palace, J., Miller, D.H., Kapoor, R. & Giovannoni, G. (2013) Biomarker report from the phase II lamotrigine trial in secondary progressive MS - neurofilament as a surrogate of disease progression. PLoS One, 8, e70019.

Golmohammadi, M., Mahmoudian, M., Hasan, E.K., Alshahrani, S.H., Romero-Parra, R.M., Malviya, J., Hjazi, A., Najm, M.A.A., Almulla, A.F., Zamanian, M.Y., Kadkhodaei, M. & Mousavi, N. (2024) Neuroprotective effects of riluzole in Alzheimer’s disease: A comprehensive review. Fundam Clin Pharmacol, 38, 225–237.

Guerreiro, R., Wojtas, A., Bras, J., Carrasquillo, M., Rogaeva, E., Majounie, E., Cruchaga, C., Sassi, C., Kauwe, J.S., Younkin, S., Hazrati, L., Collinge, J., Pocock, J., Lashley, T., Williams, J., Lambert, J.C., Amouyel, P., Goate, A., Rademakers, R., Morgan, K., Powell, J., St George-Hyslop, P., Singleton, A., Hardy, J. & Alzheimer Genetic Analysis, G. (2013) TREM2 variants in Alzheimer’s disease. N Engl J Med, 368, 117–127.

Jicha, G.A., Bowser, R., Kazam, I.G. & Davies, P. (1997) Alz-50 and MC-1, a new monoclonal antibody raised to paired helical filaments, recognize conformational epitopes on recombinant tau. J Neurosci Res, 48, 128–132.

Joel, Z., Izquierdo, P., Salih, D.A., Richardson, J.C., Cummings, D.M. & Edwards, F.A. (2018) Improving Mouse Models for Dementia. Are All the Effects in Tau Mouse Models Due to Overexpression? Cold Spring Harb Symp Quant Biol, 83, 151–161.

Jonsson, T., Stefansson, H., Steinberg, S., Jonsdottir, I., Jonsson, P.V., Snaedal, J., Bjornsson, S., Huttenlocher, J., Levey, A.I., Lah, J.J., Rujescu, D., Hampel, H., Giegling, I., Andreassen, O.A., Engedal, K., Ulstein, I., Djurovic, S., Ibrahim-Verbaas, C., Hofman, A., Ikram, M.A., van Duijn, C.M., Thorsteinsdottir, U., Kong, A. & Stefansson, K. (2013) Variant of TREM2 associated with the risk of Alzheimer’s disease. N Engl J Med, 368, 107–116.

Kapoor, R., Furby, J., Hayton, T., Smith, K.J., Altmann, D.R., Brenner, R., Chataway, J., Hughes, R.A. & Miller, D.H. (2010) Lamotrigine for neuroprotection in secondary progressive multiple sclerosis: a randomised, double-blind, placebo-controlled, parallel-group trial. Lancet Neurol, 9, 681–688.

Liu, W., Taso, O., Wang, R., Bayram, S., Graham, A.C., Garcia-Reitboeck, P., Mallach, A., Andrews, W.D., Piers, T.M., Botia, J.A., Pocock, J.M., Cummings, D.M., Hardy, J., Edwards, F.A. & Salih, D.A. (2020) Trem2 promotes anti-inflammatory responses in microglia and is suppressed under pro-inflammatory conditions. Hum Mol Genet, 29, 3224–3248.

Lo, A.C., Saab, C.Y., Black, J.A. & Waxman, S.G. (2003) Phenytoin protects spinal cord axons and preserves axonal conduction and neurological function in a model of neuroinflammation in vivo. J Neurophysiol, 90, 3566–3571.

Matarin, M., Salih, D.A., Yasvoina, M., Cummings, D.M., Guelfi, S., Liu, W., Nahaboo Solim, M.A., Moens, T.G., Paublete, R.M., Ali, S.S., Perona, M., Desai, R., Smith, K.J., Latcham, J., Fulleylove, M., Richardson, J.C., Hardy, J. & Edwards, F.A. (2015) A genome-wide gene-expression analysis and database in transgenic mice during development of amyloid or tau pathology. Cell Rep, 10, 633–644.

Morsali, D., Bechtold, D., Lee, W., Chauhdry, S., Palchaudhuri, U., Hassoon, P., Snell, D.M., Malpass, K., Piers, T., Pocock, J., Roach, A. & Smith, K.J. (2013) Safinamide and flecainide protect axons and reduce microglial activation in models of multiple sclerosis. Brain, 136, 1067–1082.

Nelson, P.T., Alafuzoff, I., Bigio, E.H., Bouras, C., Braak, H., Cairns, N.J., Castellani, R.J., Crain, B.J., Davies, P., Del Tredici, K., Duyckaerts, C., Frosch, M.P., Haroutunian, V., Hof, P.R., Hulette, C.M., Hyman, B.T., Iwatsubo, T., Jellinger, K.A., Jicha, G.A., Kovari, E., Kukull, W.A., Leverenz, J.B., Love, S., Mackenzie, I.R., Mann, D.M., Masliah, E., McKee, A.C., Montine, T.J., Morris, J.C., Schneider, J.A., Sonnen, J.A., Thal, D.R., Trojanowski, J.Q., Troncoso, J.C., Wisniewski, T., Woltjer, R.L. & Beach, T.G. (2012) Correlation of Alzheimer disease neuropathologic changes with cognitive status: a review of the literature. J Neuropathol Exp Neurol, 71, 362–381.

Potet, F., Vanoye, C.G. & George, A.L., Jr. (2016) Use-Dependent Block of Human Cardiac Sodium Channels by GS967. Mol Pharmacol, 90, 52–60.

Raftopoulos, R., Hickman, S.J., Toosy, A., Sharrack, B., Mallik, S., Paling, D., Altmann, D.R., Yiannakas, M.C., Malladi, P., Sheridan, R., Sarrigiannis, P.G., Hoggard, N., Koltzenburg, M., Gandini Wheeler-Kingshott, C.A., Schmierer, K., Giovannoni, G., Miller, D.H. & Kapoor, R. (2016) Phenytoin for neuroprotection in patients with acute optic neuritis: a randomised, placebo-controlled, phase 2 trial. Lancet Neurol, 15, 259–269.

Sadeghian, M., Mullali, G., Pocock, J.M., Piers, T., Roach, A. & Smith, K.J. (2016) Neuroprotection by safinamide in the 6-hydroxydopamine model of Parkinson’s disease. Neuropathol Appl Neurobiol, 42, 423–435.

Saito, T., Matsuba, Y., Mihira, N., Takano, J., Nilsson, P., Itohara, S., Iwata, N. & Saido, T.C. (2014) Single App knock-in mouse models of Alzheimer’s disease. Nat Neurosci, 17, 661–663.

Sanchez, P.E., Zhu, L., Verret, L., Vossel, K.A., Orr, A.G., Cirrito, J.R., Devidze, N., Ho, K., Yu, G.Q., Palop, J.J. & Mucke, L. (2012) Levetiracetam suppresses neuronal network dysfunction and reverses synaptic and cognitive deficits in an Alzheimer’s disease model. Proceedings of the National Academy of Sciences of the United States of America, 109, E2895–E2903.

Simard, J.M., Tsymbalyuk, O., Keledjian, K., Ivanov, A., Ivanova, S. & Gerzanich, V. (2012) Comparative effects of glibenclamide and riluzole in a rat model of severe cervical spinal cord injury. Exp Neurol, 233, 566–574.

Verret, L., Mann, E.O., Hang, G.B., Barth, A.M., Cobos, I., Ho, K., Devidze, N., Masliah, E., Kreitzer, A.C., Mody, I., Mucke, L. & Palop, J.J. (2012) Inhibitory interneuron deficit links altered network activity and cognitive dysfunction in Alzheimer model. Cell, 149, 708–721.

Wilson, J.R. & Fehlings, M.G. (2014) Riluzole for acute traumatic spinal cord injury: a promising neuroprotective treatment strategy. World Neurosurg, 81, 825–829.

Wood, J.I., Wong, E., Joghee, R., Balbaa, A., Vitanova, K.S., Stringer, K.M., Vanshoiack, A., Phelan, S.J., Launchbury, F., Desai, S., Tripathi, T., Hanrieder, J., Cummings, D.M., Hardy, J. & Edwards, F.A. (2022) Plaque contact and unimpaired Trem2 is required for the microglial response to amyloid pathology. Cell Rep, 41, 111686.

Yaari, Y., Selzer, M.E. & Pincus, J.H. (1986) Phenytoin: mechanisms of its anticonvulsant action. Ann Neurol, 20, 171–184.

Ziyatdinova, S., Gurevicius, K., Kutchiashvili, N., Bolkvadze, T., Nissinen, J., Tanila, H. & Pitkanen, A. (2011) Spontaneous epileptiform discharges in a mouse model of Alzheimer’s disease are suppressed by antiepileptic drugs that block sodium channels. Epilepsy Res, 94, 75–85.

